# Growth characteristics of Myco-Composites Produced Using Locally Isolated *Ganoderma Species* and Lignocellulosic Waste

**DOI:** 10.1101/2025.11.24.690336

**Authors:** Abhishek Kumar Mishra, Smruti B. Bhatt, Lakshminath Kundanati

## Abstract

This study reports the development of myco-composites from locally isolated *Ganoderma species* and regional lignocellulosic wastes such as paper, cardboard, sawdust, and wood shavings. Eight substrate formulations were evaluated for fungal colonization and composite quality. Visual inspection revealed that fine, fibrous substrates like paper and cardboard achieved the most uniform mycelial colonization and highest densities (0.45 and 0.41 g/cm^3^, respectively), while coarse, porous substrates such as wood shavings and mixed substrates (M1 and M2) produced lighter, less dense composites (0.27 g/cm^3^). Volumetric shrinkage ranged from 7.1% to 26.7%, with lower values in cardboard and higher in wood shavings. FTIR spectroscopy confirmed the presence of characteristic fungal and lignocellulosic functional groups in mixed composites. These findings demonstrate that composite density, dimensional stability, and chemical structure are strongly influenced by substrate type and proportion, highlighting the potential for tailored myco-composite materials using local waste resources.

**Graphical Abstract:** 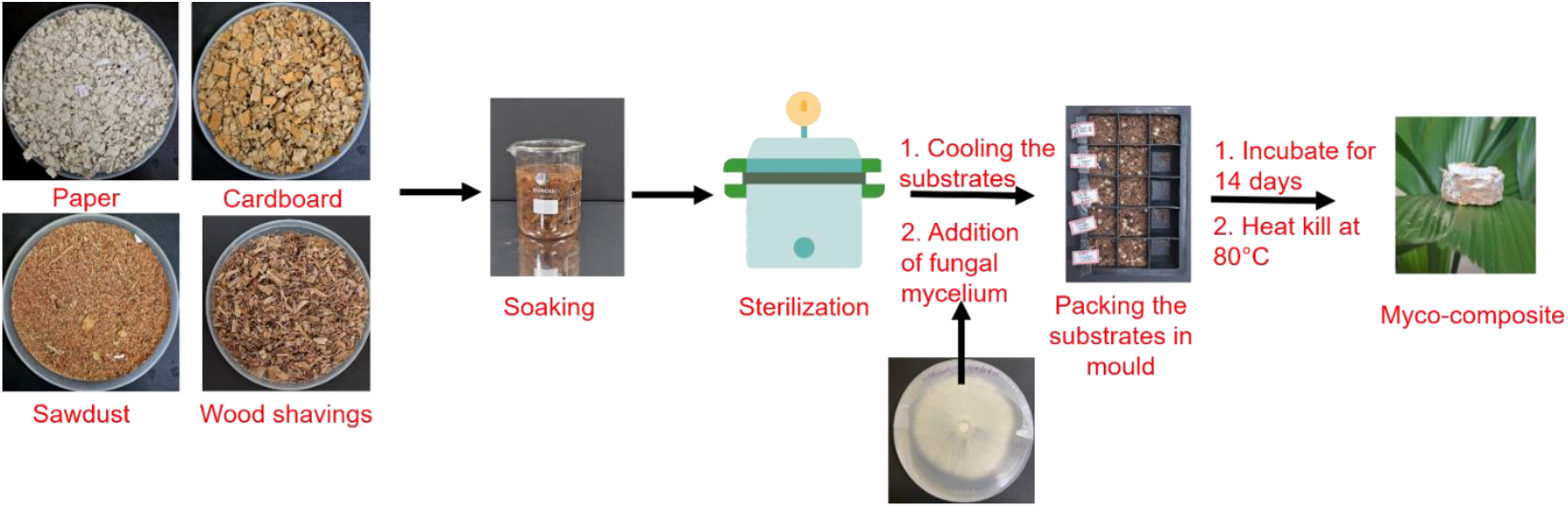

## 1. Introduction

The growing demand for sustainable and biodegradable materials has increased global interest in bio-based alternatives. Among the various bio-based alternatives, myco-composites formed via the growth of fungal mycelium on lignocellulosic substrates have gained significant attention. Myco-composites require minimal energy for production and can utilize low-cost lignocellulosic waste. Their biodegradability and low carbon footprint make them particularly suitable for packaging, insulation, and lightweight structural applications [1].

The performance of myco-composites largely depends on the fungal species used, as well as the type of the lignocellulosic substrates. *Ganoderma species* are known for forming dense, robust mycelial networks, capable of binding diverse substrates effectively. Their fast colonization, strong hyphal adhesion, and ability to grow on a wide range of lignocellulosic materials make them suitable candidates for myco-composite fabrication. Using locally isolated fungi promotes regional resource utilization and reduces dependency on commercial strains [2].

Recent studies using locally isolated fungal species have shown that wild isolates often exhibit stronger substrate specific colonization compared to commercial strains. Such species - substrate interactions strongly influence key composite parameters such as density and shrinkage, which in turn determine whether a material is better suited for secondary packaging or for insulation applications. Angelova et al., reported the isolation of *Ganoderma resinaceum* GA1M in Bulgaria and used it to develop myco-composites grown on rose flower and lavender straw substrates. The resulting materials showed densities between 0.34 and 0.46 g/cm^3^ [3].

This work aims to utilize lignocellulosic waste such as paper, cardboard, sawdust, wood shavings and locally isolated *wild Ganoderma species* for developing myco-composites. Various substrate combinations will be evaluated to assess the suitability of these abundant, non-seasonal waste as alternatives to conventional agro-waste, while enabling targeted control of composite density and shrinkage to produce materials suited for either packaging or insulation applications. The resulting myco-composites will be characterized for their physical and functional properties, including density, shrinkage, and FTIR analysis.

## 2. Materials and Method

Sawdust & wood shavings (mixture of teak and redwood) were collected from Mahalakshmi timber depot, Velachery. Paper and cardboard were collected from waste collection centre at IIT Madras campus. *Wild Ganoderma species* were isolated from IIT Madras campus. Potato dextrose agar was purchased from Hi-media, India. Jowar grains were bought from the nearby supermarket. All experiments were carried out using distilled water.

### 2.1 Growth of the *wild Ganoderma species* fungal mycelium

The fruiting body of the *wild Ganoderma species* was collected from the IIT Madras campus and tissue cultured on potato dextrose agar plates (100 mm petri-dishes), incubated at 25 °C [4]. Mycelial growth rate was calculated by measuring the radial growth extension (diameter) of mycelium on agar plates every day for a period of 15 days. It was represented in the form of milli meter/day (mm/day). Growth Area was calculated using the formula (eq. 1) given below [5].

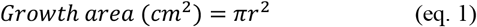

Where, r is the radius of the growth extension in cm

### 2.2 Production of mycelial spawn

The *wild Ganoderma species* mother cultures was used to prepare the mycelial spawn. To produce mycelial spawn, jowar grains were boiled in water for 30 minutes, excess moisture was removed by spreading the grains on a muslin cloth, and finally the grains were supplemented with 20 g of calcium carbonate per kilogram of grains. The treated grains were placed in glass containers and autoclaved. After cooling, 8 mm diameter mycelial agar plugs from the mother culture were inoculated into the grains to produce mycelial spawn, which was used as an inoculum to produce myco-composites [6].

### 2.3 Production of myco-composites using a combination of paper, cardboard, wood shredding’s and sawdust substrates and fungal mycelium

Four substrates namely paper, cardboard, sawdust, and wood shavings were used to prepare the myco-composites. The substrates were dried, ground, washed, and soaked for 12 hours, after which the moisture content was adjusted to 60–70%. All substrates were then sterilized in an autoclave at 121 °C and 15 psi for 20 minutes. The sterilized substrates were mixed and labelled according to the compositions provided in **Table 1** and were inoculated with 20% (w/w) mycelial spawn, ensuring uniform distribution. The inoculated mixtures were packed into sterile moulds and incubated for 14 days to allow complete colonization. After incubation, the myco-composites were dried at 80 °C for 12 hours to inactivate the fungi, and the resulting myco-composites were stored [7,8].

**Table 1.**
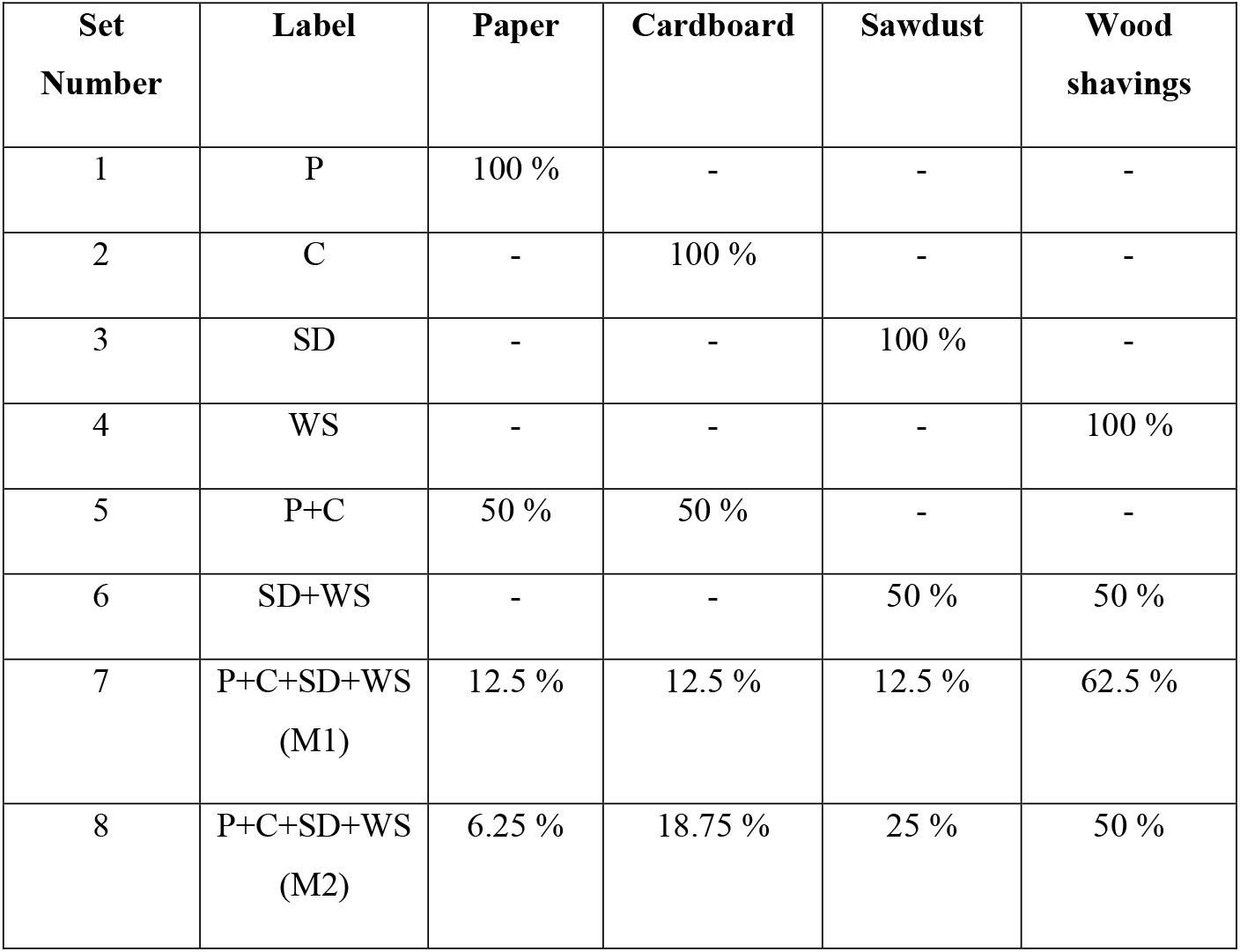
Summary of the compositions of paper, cardboard, saw-dust and wood shavings used as a substrate to produce myco-composite in this study.

### 2.4 Physical properties of the myco-composite

The density and volumetric shrinkage percentage was calculated using the following formulas (eq. 2 & 3);

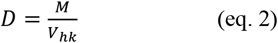

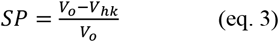

Where, D is the density of mycelial composite in g/cm^3^, M and *V*_*hk*_ are the mass and volume of dried composite respectively. SP is the shrinkage percentage and *V*_*O*_ is the volume of the mycelial composite before drying (wet) and *V*_*hk*_ is the volume of the mycelial composite after drying [5].

### 2.5 Functional properties (FTIR)

The ATR-FTIR spectra of myco-composite samples and pure mycelium sheet from agar plate was recorded within the infrared range of 4000-500 cm^−1^ in transmittance mode with 4 cm^−1^ resolution (FTIR, ATR, Perkin Elmer). Samples were dried at 50 °C to constant weight, cut into ~5–10 mm pieces, and placed directly on the ATR crystal to ensure good contact.

## 3. Results and Discussion

### 3.1 Growth of *Ganoderma species* fungal mycelium

The fruiting body of the *Ganoderma species* was collected from the IIT Madras campus **(Figure 1 a.)**. The tissue fragments taken from the underside of the fruiting body were transferred onto agar plates to get the pure fungal mycelium culture. A mycelial agar plug from the initial culture was then placed onto the centre of a fresh PDA agar plate, to monitor the radial growth for a period of 15 days **(Figure 1 b.)**. The radial mycelial growth rate of the isolated *Ganoderma species* was found to **5.84 ± 0.57 mm/day** during the linear growth phase **(Figure 1 c.)**.

**Figure 1.**
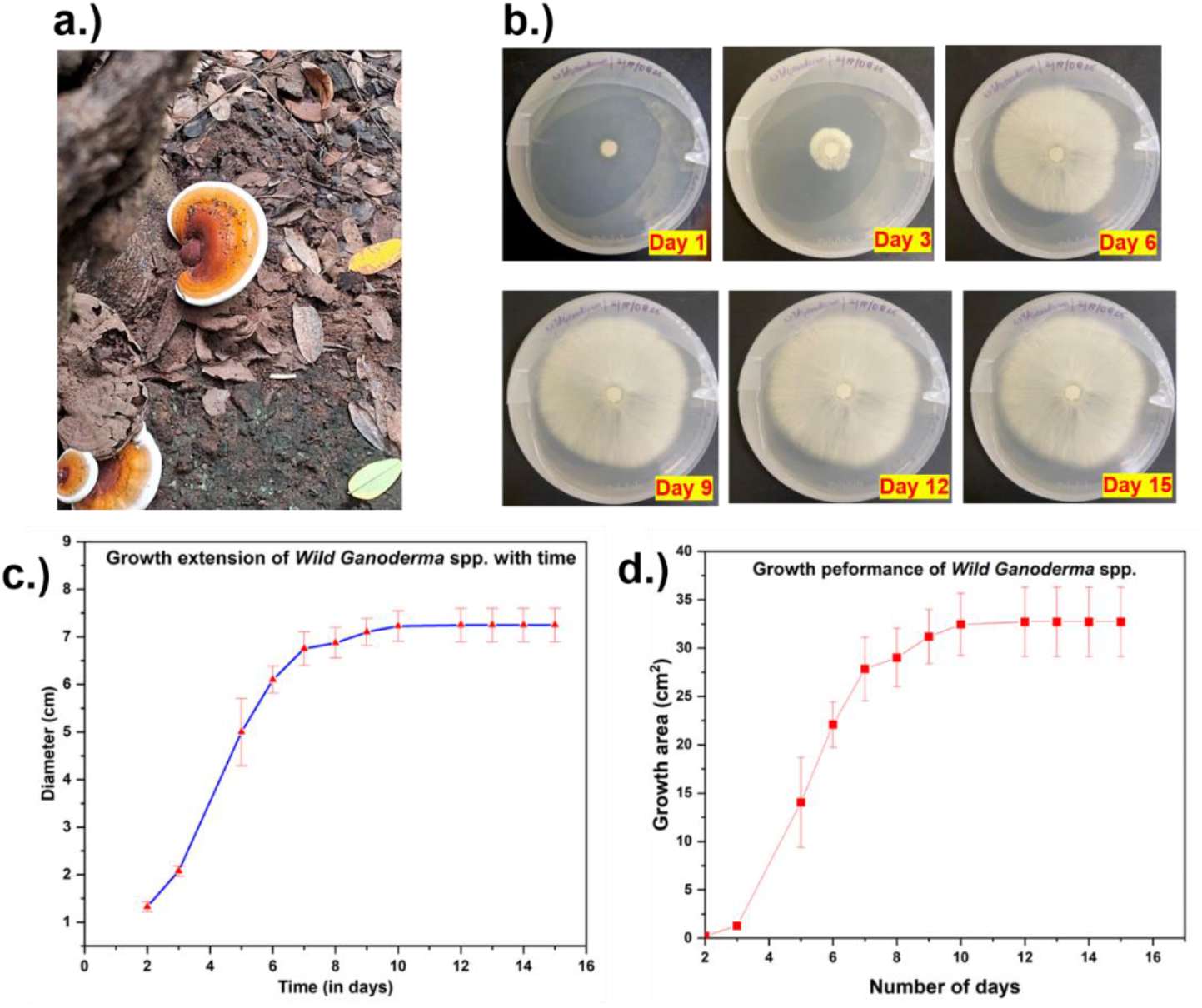
Represents a) picture of *wild Ganoderma species* isolated from IIT Madras campus, b) Progressive growth of the *wild Ganoderma species* on potato dextrose agar plates; c) Growth diameter of the *wild Ganoderma species* vs time; d) Growth area of the *wild Ganoderma species* vs time.

The growth area **(Figure 1 d.)** increased steadily from **day 2 onwards, expanding from 0.22 ± 0.08 cm**^**2**^ **to 32.4 ± 3.20 cm**^**2**^ **by Day 10**, and occupying a major portion of the 100 mm Petri dish. This growth performance indicated that the isolated fungus was a fast-colonizing strain suitable for myco-composite production. The growth rate of the *Ganoderma species* observed in this study (5.84 mm/day) was in line with the findings of Wang et al., who reported a similar radial growth rate of 6.64 mm/day for *Ganoderma lucidum* [4]. The fungal mycelial growth obtained on PDA plates was subsequently used to prepare mycelial spawn, which served as the inoculum for producing the myco-composites.

### 3.2 Visual inspection of the myco-composites

As depicted in **Figure 2**, the visual appearance of the myco-composites varied depending on the substrate used. The paper and cardboard composites appeared smooth and fully colonized, indicating good mycelial growth and strong binding. In contrast, the sawdust composite showed poor colonization and was unable to form a stable composite, resulting in low structural integrity. The wood shavings composite also showed incomplete and uneven colonization, as the large particle size of wood shavings made it difficult for the mycelium to uniformly bind the material. Among the mixed substrates, the Paper + Cardboard composite showed uniform and dense mycelial colonization, while the Sawdust + Wood shavings composite exhibited uneven growth similar to its individual components. The composites made from all four substrates (M1 and M2) showed better mycelial colonization than composites made of Saw dust /Wood Shavings, as the presence of paper and cardboard helped the mycelium spread more easily between the wood shavings/sawdust particles. However, their surfaces remained slightly rough due to the presence of these wood/sawdust fragments in the mixture.

**Figure 2.**
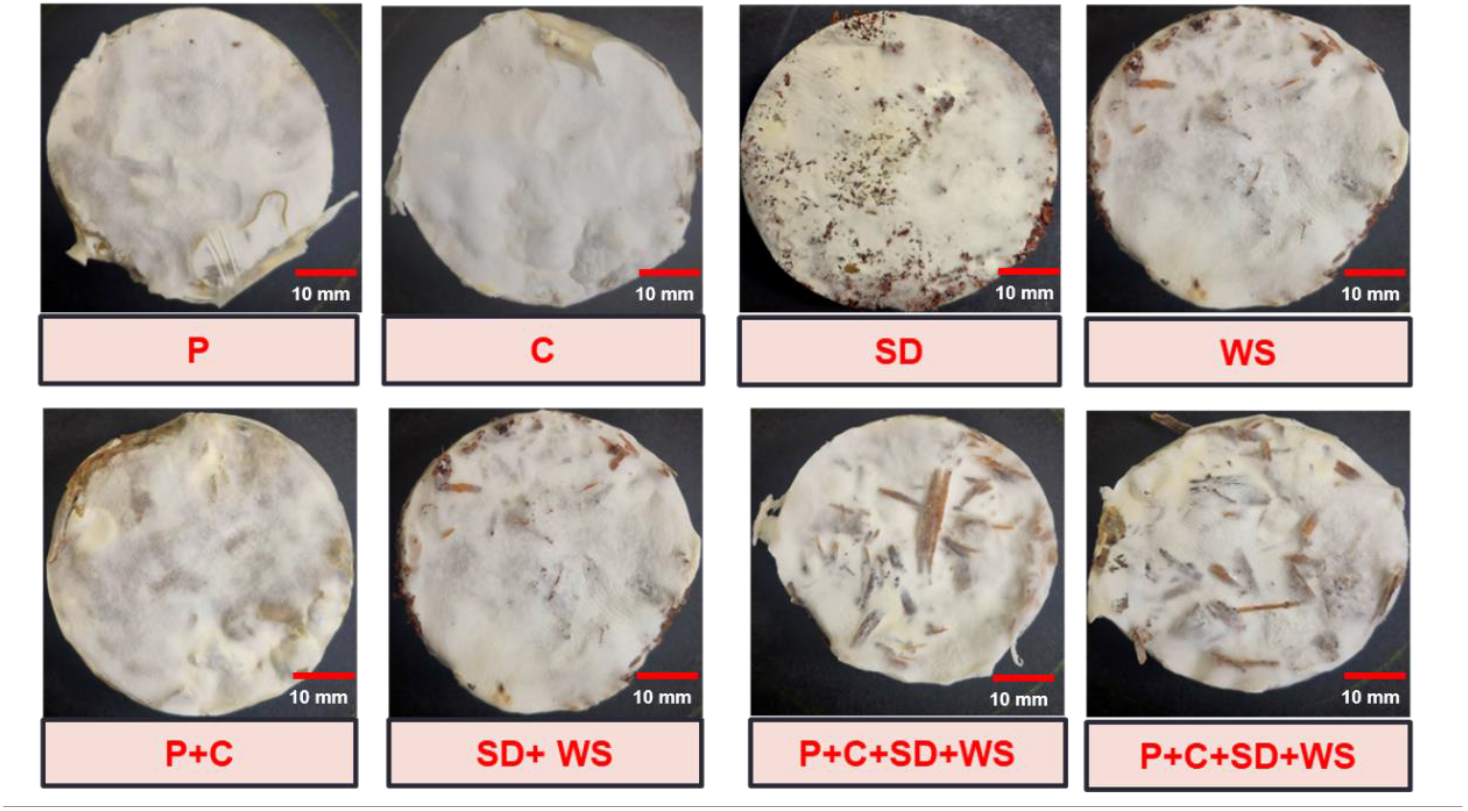
Visual images of the developed myco-composites.

### 3.3 Analysis of density and shrinkage of myco-composite

Density of the myco-composite ranged from 0.27 to 0.45 g/cm^3^ **(Figure 3 a.)**. Composites made entirely from paper (0.45 g/cm^3^) and cardboard (0.41 g/cm^3^) had the highest densities because of their fine and compressible fibres, that allowed tight packing. Composites made up of wood shavings (0.33 g/cm^3^) had lower density due to the larger structure of wood shavings, which increased porosity. Among combinations, paper + cardboard (0.40 g/cm^3^) remained dense owing to both substrates being fine and fibrous, while sawdust + wood shavings (0.39 g/cm^3^) showed moderate density because the small sawdust particles partially filled the spaces created by the larger wood shavings. The mixed substrates M1 (0.27 g/cm^3^) and M2 (0.32 g/cm^3^) recorded the lowest densities, primarily because the higher proportion of wood shavings in both mixtures led to a more open and porous internal structure. Overall, materials with fine, fibrous texture produced denser composites, while those containing larger particles resulted in lighter, more porous structures. The density of the myco-composites developed in this study (0.27-0.45 g/cm^3^) was in line with the values reported by Appels et al., which ranged from 0.10 to 0.39 g/cm^3^ [9]. The percentage volumetric shrinkage of the myco-composite ranged from 7.07 to 26.65 % **(Figure 3 b.)**. The volumetric shrinkage of the composites varied with substrate type, with the highest values in sawdust (26.65%) and sawdust + wood shavings (23.94%) due to their loose, moisture-retaining structure. Cardboard showed the lowest volumetric shrinkage (7.07%), followed by paper (17.87%) and paper + cardboard (17.20%), indicating that finer, denser fibres help maintain dimensional stability. Composites made up of all the substrates - M1 and M2 (23.24%, 22.56%) showed higher volumetric shrinkage because of the larger proportion of wood shavings (50-62.5%).

**Figure 3.**
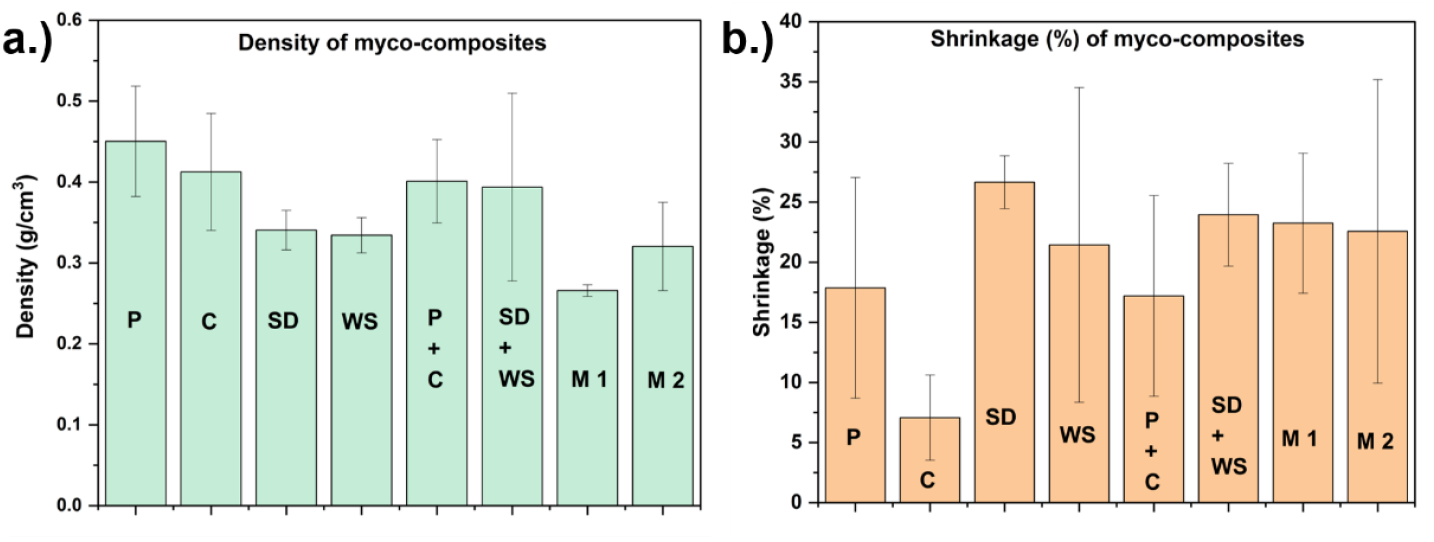
Density (a) and Volumetric Shrinkage (b) of myco-composites (P-paper, C-Cardboard, SD-Sawdust; WS-Wood Shaving’s).

### 3.4 Functional property of the myco-composite

The FTIR analysis of the pure mycelium sheet and M1 composite (Paper +Cardboard +Sawdust+ Wood Shavings) highlights differences in chemical bonding and functional groups present in each material **(Figure 4)**. The pure mycelium sheet shows prominent peaks at 3274 cm^−1^ (O-H stretching, indicating polysaccharides), 2922 cm^−1^ (C-H stretching from aliphatic chains), 1626 cm^−1^ and 1537 cm^−1^ (amide I and II bands, characteristic of proteins), 1390 cm^−1^ (C-H bending), and 1030 cm^−1^ (C-O stretching from polysaccharides) [10,11]. In the M1 composite, peaks appear at 3312 cm^−1^ (broad peak, O-H stretching vibration), 2922 cm^−1^ (C-H stretching), 1638 cm^−1^ (stretching frequency of C=O of amide group, indicating protein), and 1030 cm^−1^ (C-O stretching from polysaccharides) [11,12].

**Figure 4.**
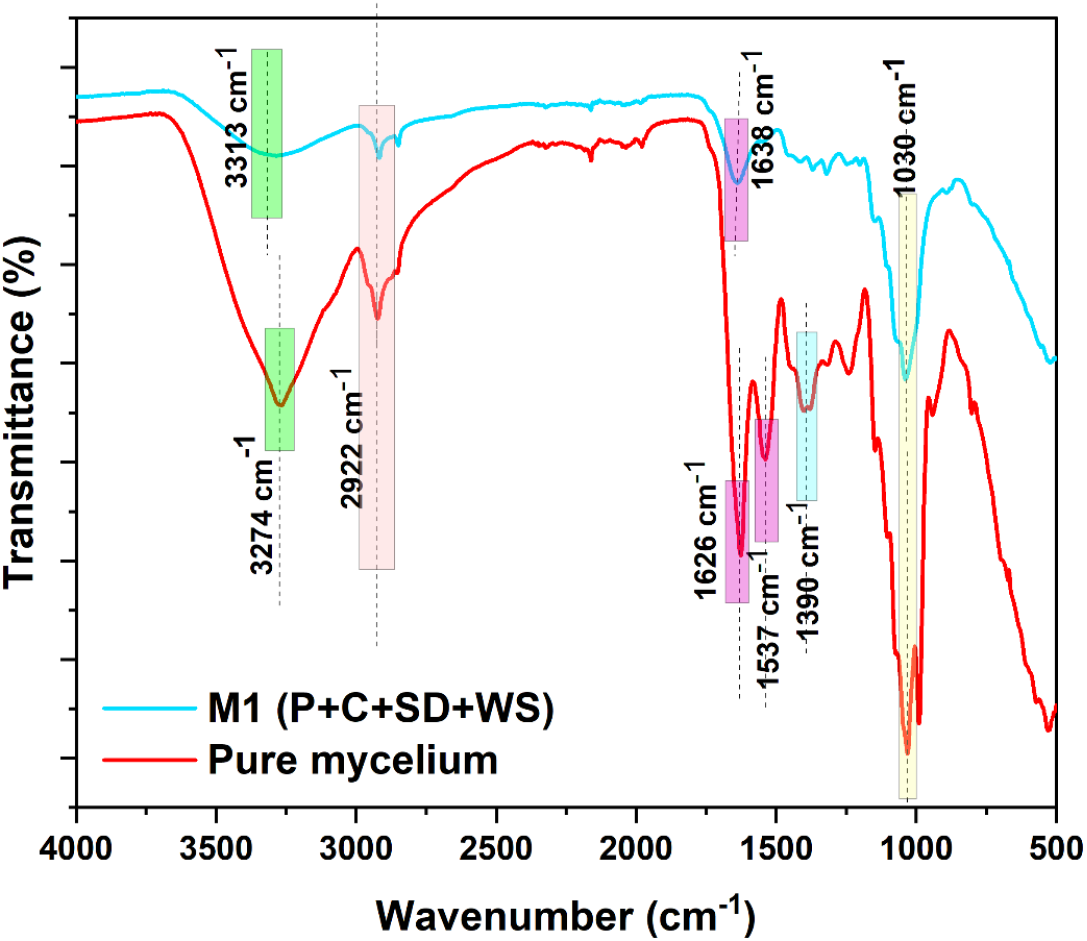
FTIR spectra of myco-composite (M1) and pure mycelium sheet from agar plate.

The FTIR profiles of the pure mycelium sheet and the M1 composite showed similar characteristic peaks, confirming successful fungal colonization of the composite. The presence of the amide band in both the pure mycelium sheet and M1 composite suggests molecular interactions arising from the growth of mycelium on the lignocellulosic substrates, where fungal cell-wall proteins and polysaccharides interact with lignocellulosic components during colonization [13]. A similar result was described by Aguilar et al., who observed a distinct amide band at approximately 1620 cm^−1^ in the FTIR spectra of both the myco-composite and the fungal liquid culture [13]. Parhizi et al., similarly reported a peak at 1650 cm^-1^ corresponding to amide groups in both pure mycelium and myco-composites [14].

## 4. Conclusion

In this study, the type of lignocellulosic substrate was shown to strongly influence the growth, density and shrinkage of myco-composites produced using locally isolated *Ganoderma species*. Fine, fibrous substrates (paper, cardboard) produced dense, stable composites, while coarse substrates (wood shavings) yielded lighter, porous composites. FTIR analysis of the composites revealed the presence of both fungal and lignocellulosic functional groups, indicating interaction between mycelium and substrate. Overall, the findings highlight the potential of these myco-composites to serve as alternatives for packaging, insulation, and other environmentally conscious applications.

## References

[1] W. Aiduang, P. Jinanukul, W. Thamjaree, T. Kiatsiriroat, T. Waroonkun, S. Lumyong, A Comprehensive Review on Studying and Developing Guidelines to Standardize the Inspection of Properties and Production Methods for Mycelium-Bound Composites in Bio-Based Building Material Applications, Biomimetics 9 (2024) 549. 10.3390/biomimetics9090549.

[2] K.K. Alaneme, J.U. Anaele, T.M. Oke, S.A. Kareem, M. Adediran, O.A. Ajibuwa, Y.O. Anabaranze, Mycelium based composites: A review of their bio-fabrication procedures, material properties and potential for green building and construction applications, Alexandria Engineering Journal 83 (2023) 234–250. 10.1016/j.aej.2023.10.012.

[3] G. Angelova, M. Brazkova, P. Stefanova, D. Blazheva, V. Vladev, N. Petkova, A. Slavov, P. Denev, D. Karashanova, R. Zaharieva, A. Enev, A. Krastanov, Waste Rose Flower and Lavender Straw Biomass—An Innovative Lignocellulose Feedstock for Mycelium Bio-Materials Development Using Newly Isolated Ganoderma resinaceum GA1M, Journal of Fungi 7 (2021) 866. 10.3390/jof7100866.

[4] J.G. van den Brandhof, N. Hansen, C. Hou, S.C. Broers, M. Tegelaar, H.A.B. Wösten, Characterization of pure mycelium materials from different mushroom-forming fungi, Antonie van Leeuwenhoek 118 (2025) 121. 10.1007/s10482-025-02133-5.

[5] Y. Wang, G. Hausner, P.R. Rout, Q. Yuan, Investigation of fungal mycelium-bound bio-foams from agricultural wastes as sustainable and eco-conscious packaging innovations, Journal of Cleaner Production 501 (2025) 145206. 10.1016/j.jclepro.2025.145206.

[6] S. Subedi, N. Kunwar, K.R. Pandey, Y.R. Joshi, Performance of oyster mushroom (Pleurotus ostreatus) on paddy straw, water hyacinth and their combinations, Heliyon 9 (2023) e19051. 10.1016/j.heliyon.2023.e19051.

[7] E. Elsacker, S. Vandelook, J. Brancart, E. Peeters, L.D. Laet, Mechanical, physical and chemical characterisation of mycelium-based composites with different types of lignocellulosic substrates, PLOS ONE 14 (2019) e0213954. 10.1371/journal.pone.0213954.

[8] E. Elsacker, S. Vandelook, A. Van Wylick, J. Ruytinx, L. De Laet, E. Peeters, A comprehensive framework for the production of mycelium-based lignocellulosic composites, Science of The Total Environment 725 (2020) 138431. 10.1016/j.scitotenv.2020.138431.

[9] F.V.W. Appels, S. Camere, M. Montalti, E. Karana, K.M.B. Jansen, J. Dijksterhuis, P. Krijgsheld, H.A.B. Wösten, Fabrication factors influencing mechanical, moisture- and water-related properties of mycelium-based composites, Materials & Design 161 (2019) 64–71. 10.1016/j.matdes.2018.11.027.

[10] V. Kachrimanidou, A. Papadaki, H. Papapostolou, M. Alexandri, Z. Gonou-Zagou, N. Kopsahelis, Ganoderma lucidum Mycelia Mass and Bioactive Compounds Production through Grape Pomace and Cheese Whey Valorization, Molecules 28 (2023) 6331. 10.3390/molecules28176331.

[11] G.S. Cano-Díaz, A. Rosas-Aburto, E. Vivaldo-Lima, L. Flores-Santos, M.A. Vega-Hernández, M.G. Hernández-Luna, A. Martinez, Determination of the Composition of Lignocellulosic Biomasses from Combined Analyses of Thermal, Spectroscopic, and Wet Chemical Methods, Ind. Eng. Chem. Res. 60 (2021) 3502–3515. 10.1021/acs.iecr.0c05243.

[12] M. Shi, Z. Zhang, Y. Yang, Antioxidant and immunoregulatory activity of Ganoderma lucidum polysaccharide (GLP), Carbohydrate Polymers 95 (2013) 200–206. 10.1016/j.carbpol.2013.02.081.

[13] K. Aguilar, L. Figel, S. Saker, L. Soufflet, N. Brosse, A. Besserer, Steam Explosion: A Booster for Fungal Growth in a Myco-composite, ACS Sustainable Chem. Eng. 12 (2024) 11650–11659. 10.1021/acssuschemeng.4c03099.

[14] Z. Parhizi, J. Dearnaley, K. Kauter, D. Mikkelsen, P. Pal, T. Shelley, P. (Polly) Burey, The Fungus Among Us: Innovations and Applications of Mycelium-Based Composites, Journal of Fungi 11 (2025) 549. 10.3390/jof11080549.

